# Understanding *Campylobacter coli* isolates from the Vietnamese meat production network; a pilot study

**DOI:** 10.1101/2023.11.10.566519

**Authors:** Burhan Lehri, Georgina Navoly, Abigail Corser, Fauzy Nashar, Sam Willcocks, Pham Thi Ngoc, Brendan W. Wren, Luu Quynh Huong, Richard A. Stabler

## Abstract

Changing farming practices and the associated increase in the use of antibiotics are amongst the main drivers shaping the global increase of Campylobacter infections. The effects farming practices have on *Campylobacter* species, need to be studied at the global scale, particularly in emerging middle-income countries, where the demand for low-cost poultry meat is rising. While *C. jejuni* causes the majority of poultry associated diarrhoea, *C. coli* causes a significant amount of disease but are relatively understudied. In this study we characterised seven *C. coli* strains isolated from poultry farms and markets in Hanoi, Vietnam. Comprehensive data sets of bacterial Whole-Genome Sequencing; and phenotypic assays, such as, growth, motility, antimicrobial resistant testing along with virulence testing were performed to reveal the genetic relatedness and pathophysiological characteristics of seven *C. coli* strains. Six isolates were classified as multi-drug resistant, with all isolates resistant to ciprofloxacin, nalidixic acid and tetracycline, but susceptible to phenicols. All isolates had similar growth rates, while five were hyper-motile. Lethality of the isolates towards a tractable host-model system, larvae of the greater wax moth *Galleria mellonella*, often used to determine *Campylobacter* virulence was demonstrated for the first time for *C. coli*. Multilocus sequence typing data correlates with North American, European, and Asian isolates from patients suffering from gastroenteritis, emphasising the global spread of these strains. This work demonstrates that *C. coli*, with high levels of antimicrobial resistance, is an understudied global threat.

**Data summary:** GenBank database with accession numbers JAKGTW000000000, JAKGTV000000000, JAKGTS000000000, JAKGTU000000000, JAKGTT000000000, JAKGTR000000000 and CP091310

https://www.ncbi.nlm.nih.gov/nuccore/JAKGTW000000000

https://www.ncbi.nlm.nih.gov/nuccore/JAKGTV000000000

https://www.ncbi.nlm.nih.gov/nuccore/JAKGTS000000000

https://www.ncbi.nlm.nih.gov/nuccore/JAKGTU000000000

https://www.ncbi.nlm.nih.gov/nuccore/JAKGTT000000000

https://www.ncbi.nlm.nih.gov/nuccore/JAKGTR000000000

https://www.ncbi.nlm.nih.gov/nuccore/CP091310.1

The authors confirm all supporting data, code and protocols have been provided within the article or through supplementary data files.

## 1. Introduction

The demand for low-cost poultry meat and eggs are rapidly rising globally. To satisfy the increasing demand for meat and egg production the use of antimicrobial agents in animal food production has become widespread (Mulchandani et al. 2023). The global use of antibiotics is also projected to rise (Van Boeckel et al. 2017). Farmers, particularly in rapidly developing countries often rely on antibiotics as growth promoters and to improve the health of animals, however, this largely contributes to the generation of antibiotic resistant bacteria and the subsequent global increase in antimicrobial resistance (AMR) (Agyare et al. 2018). The seriousness of antibiotic overuse has been exacerbated further by difficulties in implementing antibiotic regulation policies in several countries (WHO 2017, Tangcharoensathien et al. 2018). The interplay between the increasing incidence of foodborne illnesses, bacterial infections and AMR is strongly suggesting that the expansion of poultry production is presenting a growing public health risk (Kaakoush et al. 2015, Skarp et al. 2016).

The rapid intensification of poultry and egg has been linked with increasing incidence of bacterial food poisoning, mostly caused by *Campylobacter* species. Campylobacter spp are estimated to cause globally approximately 166M cases of disease (92M-300M 95% UI (uncertainty intervals)) and 37K (27-55k 95% UI) deaths annually (Kirk et al. 2015). This equates to 1,390 (752-2,576 95% UI) Disability Adjusted Life Years (DALYs) per 100,000 persons and 31 (22-46 UI) deaths per 100,000 persons (Kirk et al. 2015). However, LMIC regions carry significantly more burden of disease with Campylobacter spp being the most prevalent foodborne disease in the WHO South-East Asia region (1,152 [200-3,372 95% UI] DALYS per 100,000 persons and 0.4 [0.1-0.9 95% UI] deaths per 100,000 persons (Kirk et al. 2015). While species of Campylobacter are often combined in epidemiological data *C. jejuni* receives the majority of study. *C. coli*, the second most common *Campylobacter* species, is also implicated in bacterial gastroenteritis, contributing to a quarter of *Campylobacter* related gastroenteritis (Kaakoush et al. 2015), and one of the leading cause of foodborne diarrhoea (Laaveri et al. 2018).

Vietnam has undergone rapid economic and population growth, along with urbanisation leading to changes in agricultural and food production practices. Poultry is most popular livestock in Vietnam and the second-largest producer of meat (Birhanu et al. 2021). Vietnam had 512.7 million birds in 2020 producing 1.505 million tonnes of meat and 15.1 billion eggs (Birhanu et al. 2021). There is a paucity of data on Campylobacter prevalence and epidemiological data in Vietnam, in particularly for *C. coli*. A previous Vietnamese study conducted, identified *C. coli* being more resistant to antibiotics, such as ciprofloxacin, nalidixic acid and tetracycline, than *C. jejuni*, even though both species belong to the same ecological niche (Nguyen et al. 2016).

A survey of 30 chicken farms and 30 retail markets in Thai Nguyen province, Vietnam. A total of 36 putative Campylobacter isolates were isolated from farms 25/30 (83,3%) and from market 11/30 (36,7%) following the ISO 10272-1: 2017. The species identification using multiplex PCR identified *C. jejuni* (29/39, 80,6%) followed by *C. coli* (7/36, 19,4%). The aim of this pilot study was to isolate and characterise understudied *C. coli* isolated from poultry production networks, sampling originating from farms and point-of-sale retail outlets in Hanoi, Vietnam. Each isolate were to be investigated for phenotypic AMR and virulence traits and compared to each isolates draft genome. Additionally, the *Galleria mellonella* virulence infection model was adapted for *C. coli*, which has not been previously reported. This pilot study paves the way for the large in-depth One Health Poultry Hub investigation of Campylobacter across Vietnam and provides much needed data on *C. coli* present in the Vietnamese poultry meat production network.

## 2. Material and methods

### 2.1 Growth conditions

*C. coli* isolates were stored at -80 °C and grown for 48 hours on Columbia Blood Agar (CBA) base (Oxoid, United Kingdom) supplemented with 5% defibrinated horse blood (Oxoid, UK) and Skirrow (*Campylobacter* selective supplement, Oxoid, UK). Isolates were grown under microaerobic conditions (10% CO_2_, 5% O_2_, and 85% N_2_) in a variable atmospheric incubator (Don Whitley, Scientific, UK) at 37 °C. *C. coli* isolates were re-streaked for another 24 hours on CBA/blood/skirrow plates prior to experimentation. Oxidase, indole test and gram staining were used to confirm the presence of *Campylobacter* sp. (Table A1). Hippurate test was used to distinguish *C. jejuni* (positive) from *C. coli* (negative) (Table A1). Growth curve experiments were also conducted to compare *C. coli* growth rate with well-studied *C. jejuni* 81-176. Growth curves were averaged from two biological repeats, in 10 ml Brucella broth (Oxoid, UK), containing ∼10^8^ colony-forming units (CFU), grown for 24 hours under microaerobic conditions (10% CO_2_, 5% O_2_, and 85% N_2_) at 37 °C with optical density readings (OD_600nm_), being taken at 2-hour intervals.

### 2.2 Motility assay

Overnight *C. coli* cultures were diluted to an OD_600nm_ of 0.2 in Brucella broth (Oxoid, UK). *C. coli* (3 µl; 8 x 10^8^ CFU/ml bacteria) was added into 0.4% (wt/vol) Brucella agar and incubated for 72 hours at 37 °C, under microaerobic conditions. Motility measurements were recorded 24, 48 and 72 hours. For experiment controls the representative strains *C. jejuni* 81-176 (a human isolate with a putative plasmid containing a type IV secretion and verified as highly virulent through testing in macaque monkeys (Russell et al. 1989) and *C. jejuni* 11168H (a hyper-motile variant of the standard original sequenced strain *C. jejuni* NCTC11168 (Parkhill et al. 2000)) were used. *C. jejuni* strains were used as they have well characterised motility and allow relative motility between *C. coli* and *C. jejuni*.

### 2.3 Antimicrobial susceptibility testing

Minimum inhibitory concentration (MIC) was determined based on EUCAST, ISO 20776-1:2019 and CLSI guidelines, following the broth microdilution method in untreated 96 well Nunc Sterile round bottom plates (ThermoFisher Scientific, UK). 5 x 10^5^ CFU/ml of overnight bacterial cultures were inoculated with varying concentrations of antibiotics, described below. Antibiotics were prepared in correlation with EUCAST, ISO 20776-1:2019 guidelines. The samples were incubated for 48 hours at 37 °C in filter sterilised cation adjusted MH broth, supplemented with 2.5% defibrinated horse blood (Oxoid, UK) along with 20 mg/L β- Nicotinamide adenine dinucleotide sodium salt (Sigma Aldrich, UK), at 120 revolutions per minute (RPM), under microaerobic conditions. Growth was determined visually by comparing to the negative control, as well as by using a SpectraMax iD5 microplate reader (Molecular Devices, UK).

Antibiotics tested with ranges and breakpoints as described by National Antimicrobial Resistance Monitoring System for Enteric Bacteria (NARMS) (https://www.cdc.gov/narms/antibiotics-tested.html) unless stated. Tentative epidemiological cut-off values (T)ECOFF were used for streptomycin [4 mg/L], ampicillin [16 mg/L], as no breakpoints were available during the study (EUCAST 2020) however kanamycin had no breakpoint or T(E)ECOFF values available. Briefly Aminoglycosides; gentamicin (0.125-32 mg/L, S ≤ 2 mg/L, R ≥ 4 mg/L), kanamycin (0.5-16 mg/L, no break points), streptomycin (0.0625-64 mg/L, S ≤ 4 mg/L, R ≥ 8 mg/L). Glycosamides; clindamycin (0.0156-32 mg/L, S ≤ 1 mg/L, R ≥ 2 mg/L). Macrolide; azithromycin (0.015-64 mg/L, S ≤ 0.5 mg/L, R ≥ 1 mg/L), erythromycin (0.25-64 mg/L, S ≤ 8 mg/L, R > 8 (EUCAST 2023)). Phenicol; chloramphenicol (0.0156-256 mg/L, S ≤ 16 mg/L, R ≥ 32 mg/L), florfenicol (0.03-64 mg/L, S ≤ 4 mg/L, R ≥ 8 mg/L). Quinolone: ciprofloxacin (0.0156-64 mg/L, S ≤ 0.001, R > 0.5 (EUCAST 2023)), nalidixic acid (2-64 mg/L, S ≤ 16 mg/L, R ≥ 32 mg/L). Tetracycline (0.0625-64 mg/L, S ≤ 2 mg/L, R ≥ 4 mg/L). Ketolide; telithromycin (0.0156-8 mg/L, S ≤ 4 mg/L, R ≥ 8 mg/L). β- lactam; ampicillin (0.015-125 mg/L, S ≤ 16 mg/L, R ≥ 32 mg/L). *C. jejuni* ATCC 33560 with known MIC values was used as a control (EUCAST, ISO 20776-1:2019).

### 2.4 Virulence testing using *Galleria mellonella* model

*Campylobacter* isolates were measured to an OD_600nm_ of 0.1 (corresponding to 1 x 10^8^ CFU/ml bacteria) in Phosphate-buffered saline (PBS), 10 µl was injected into *G. mellonella* larvae (Livefoods Direct Ltd, UK) foreleg. *G. mellonella* was incubated at 37 °C with phenotypic effect being monitored over the course of nine days. Three biological replicates (using 2 x n=5 *G. mellonella*, 1 x n=10 *G. mellonella*) were conducted over 9 days. Phosphate-buffered saline (PBS) was used as a negative control and virulent *C. jejuni* 81-176 was used as a positive control. *C. jejuni* 81-176 was used as a control as the species has previously been used for the *G. mellonella* model (Champion et al. 2010, Senior et al. 2011) while, to our knowledge no *C. coli* strains have previously been studied in the *G. mellonella* model.

### 2.5 Genomic DNA extraction and sequencing

Genomic DNA was extracted using PureLink™ Genomic DNA Mini Kit (Invitrogen, UK), following manufacturer guidelines. Prior to sequencing DNA quality and concentration was measured using a NanoDrop 2000/2000c Spectrophotometer (Thermofisher Scientific, UK). Illumina Miseq library preparation and sequencing were prepared according to manufacturer’s protocol. Libraries were sequenced on a MiSeq (Illumina, USA) with 2 x 150 bp reagent kit v2. Additional sequencing was conducted using a MinION sequencer (Oxford Nanopore, UK). A R10 flow cell (FLO-MIN111) was used for sequencing, alongside ligation sequencing kit (SQK-LSK110) and the rapid barcoding kit (SQK-RBK004) for multiplexing, manufacturer recommended PCR conditions were used with eight minutes amplification using NEB LongAmp Taq 2X Master Mix (NEB M0287). DNA end repair was performed using NEBNext Ultra II End repair/dA-tailing Module (E7546), while adapter ligation was done using the NEBNext Quick Ligation Module (E6056).

### 2.6 Genome assembly

Raw fastq files were quality controlled using TrimmomaticPE with settings: LEADING:3 TRAILING:3 SLIDINGWINDOW:4:20 MINLEN:36 (Bolger et al. 2014). Sequenced reads were *de novo* assembled using St. Petersburg genome assembler (SPAdes version 3.15.3) (Nurk et al. 2013). The draft genome assemblies were assessed using quality assessment tool for genome assemblies (Quast) (Gurevich et al. 2013). For *C. coli* isolates that were sequenced on both MinION and MiSeq, draft genomes were assembled using Unicycler v0.4.9 (Wick et al. 2017). Once assembled, contigs were annotated using Prokaryotic Genome Annotation (Prokka) (Seemann 2014) using a bespoke *Campylobacter* database.

### 2.7 AMR gene identification, MLST and virulence analysis

Abricate with the ResFinder database (update date 10-03-2022) (Zankari et al. 2012, Seemann 2018) was used to identify AMR genes. Additionally, PathogenWatch (https://pathogen.watch/) identified AMR related single point mutations within the *C. coli* genomes. Multi-Locus Sequence Typing (MLST) analysis was conducted with MLST 2.0 from the Centre for Genomic Epidemiology using PubMLST.org database version 2.0.0. PubMLST database was also used to identify clonal complex of the sequenced isolates. ChewBBACA (version 16.04) (Silva et al. 2018) with the *C. jejuni* schema ‘INNUENDO_wgMLST’ (update date 20-05-2021) having 2794 loci was used for core genome multi locus sequence typing (cgMLST), by selecting loci that are present in 95 % of *C. coli* and *C. jejuni* isolates within RefSeq database. PHYLOViZ was used to generate a minimum spanning tree, the Newick tree format, which was imported to Interactive tree of life (iTOL) to generate the tree in Figure 4. For ChewBBACA and Abricate *C. coli* isolates were compared against complete RefSeq *C. jejuni* and *C. coli* isolates. Pre-annotated sequence files were imported onto the *Campylobacter* Virulence Factors of Pathogenic Bacteria database (VFDB) (http://www.mgc.ac.cn/cgi-bin/VFs/genus.cgi?Genus=Campylobacter).

### 2.8 Statistical analyses

Survival curves were generated using Prism version 9.1 software, and survival rates of *G. mellonella* models were analysed using the Kaplan-Meier method. Tests were performed using a two-sided probability, and survival rates were considered statistically significant when p < 0.05. For motility and growth rate analysis, three biological replicates were conducted, and statistical significance was conducted using one way ANOVA test, with p < 0.05 being considered statistically significant.

## 3. Results

### 3.1 Phenotypic analysis

#### 3.1.1 Growth rate analysis of *C. coli*

*C. coli* were isolated from poultry faeces (C57, C75, C76) and meat (M8, M10, M14, M39) from farms, retail markets or supermarkets in Hanoi, Vietnam (Supplementary Table A1). All isolates were catalase and oxidase positive but Indole and Hippurate negative (Supplementary Table A1). *C. coli* growth was also compared with *C. jejuni* 81-176 in brucella broth for 24 hours, demonstrating no significant difference in growth rate between the *C. coli* isolates and *C. jejuni* 81-176 (Supplementary Figure S1).

#### 3.1.2 Motility

To assess motility of the *C. coli* strains, the isolates were compared to two *C. jejuni* with known motility phenotypes; hyper-motile *C. jejuni* 11168H (Karlyshev et al. 2001, Macdonald et al. 2017) and motile *C. jejuni* 81-176 (Korlath et al. 1985, Hofreuter et al. 2006). *C. coli* isolates C75 (p=0.001), M14 (p=0.01) and M39 (p = 0.04) were more motile, than *C. jejuni* 11168H; (Figure 1). *C. coli* C57 (p=0.15) and C76 (p=0.06) motility was the same as *C. jejuni* 11168H (Figure 1). *C. coli* M8 and M10 were as motile as *C. jejuni* 81-176; but significantly less motile than *C. jejuni* 11168H, p=0.02 and 0.03, respectively (Figure 1).

**Figure 1.**
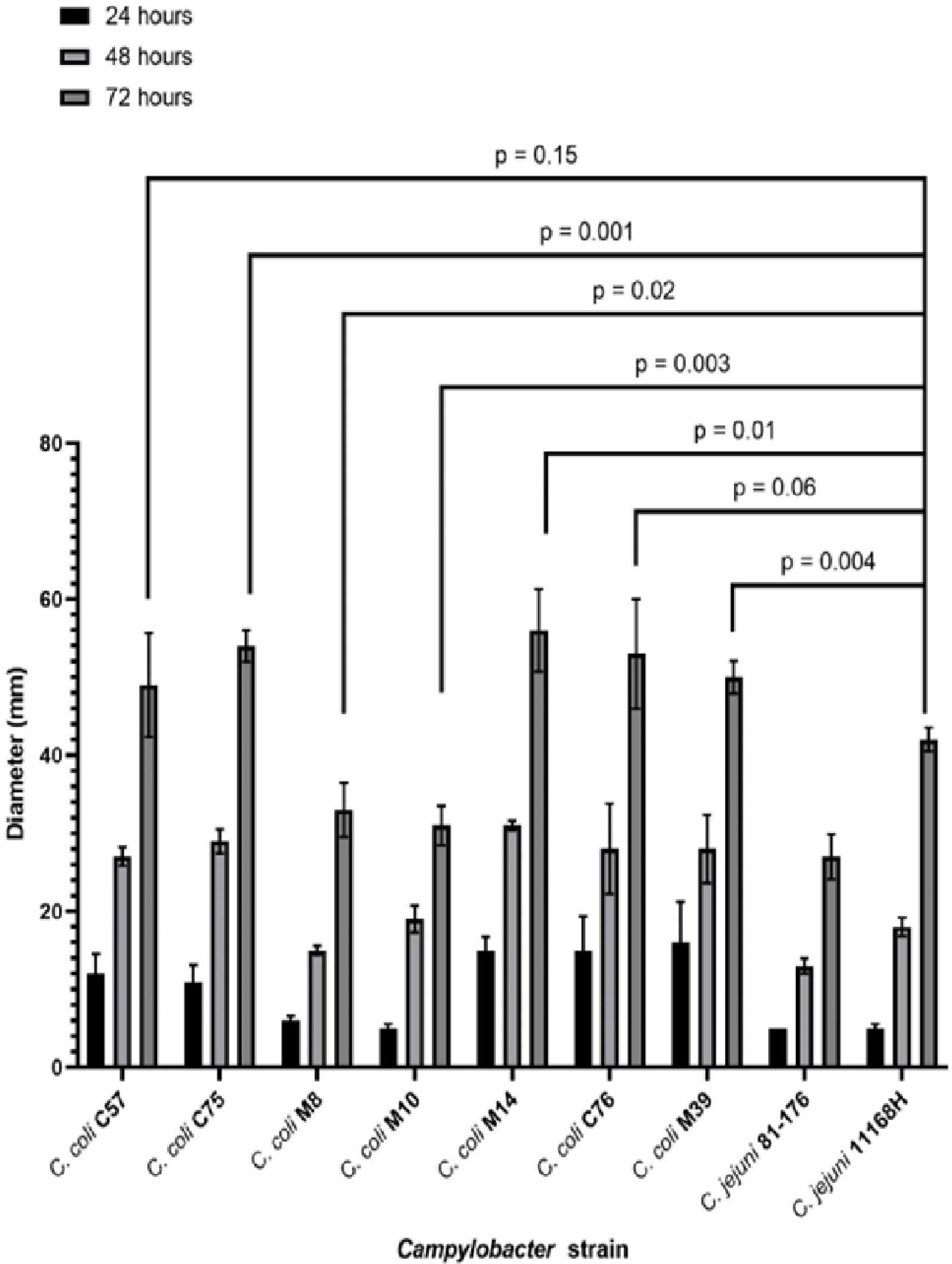
Motility analysis of *C. coli* isolates along with *C. jejuni* 81-176 and 11168H. The data shows an average of three biological replicates. Data was obtained by inoculating Brucella 0.4% (wt/vol) agar with 8 x 10^8^ CFU/ml bacteria. Growth was observed over three days with measurement time points being 24, 48 and 72 hrs. P values are generated using one-way ANOVA. *C. coli* isolates were compared against *C. jejuni* 81-176 and hyper-motile isolate *C. jejuni* 11168H.

#### 3.1.3 Antibacterial sensitivity testing

All *C. coli* isolates were resistant to quinolones (ciprofloxacin/nalidixic acid) and tetracycline (Figure 2, Table 1). For aminoglycosides, *C. coli* C57 and C76 were resistant to gentamycin. C57 and C76 also had kanamycin MIC of >16 mg/L and streptomycin MIC of 64 mg/L, which was greater than the (T)ECOFF of 4 mg/L (Figure 2, Table 1). All isolates were sensitive to phenicols, however, C75 and M8 were a single doubling dilution below the resistance cut-off (Figure 2, Table 1). *C. coli* C57, C76 and M8 were resistant to macrolides (erythromycin and azithromycin) and lincosamide (clindamycin) (Figure 2, Table A2). Isolates C57 and M8 were also resistant to telithromycin (Figure 2, Table 1). When comparing β-Lactam (ampicillin) resistant data isolates; C57, M39 and C75 had a MIC of 125 mg/L, significantly higher than (T)ECOFF (16 mg/L), while *C. coli* M8 (1.9 mg/L) and M14 (7.8125 mg/L) had MIC lower than (T)ECOFF. Multi-drug resistance (MDR) of thermophilic *Campylobacter* strains is defined as resistance to at least three different antimicrobial classes (Magiorakos et al. 2012). All isolates, except *C. coli* M14, match the MDR definition. *C. coli* M14 was only resistant to quinolones and tetracycline, however, was also above the (T)ECOFF for streptomycin (aminoglycoside; (16 vs 4 mg/L) (Table 1). *C. coli* C57 was resistant to all classes tested, except phenicols (Table 1).

**Figure 2.**
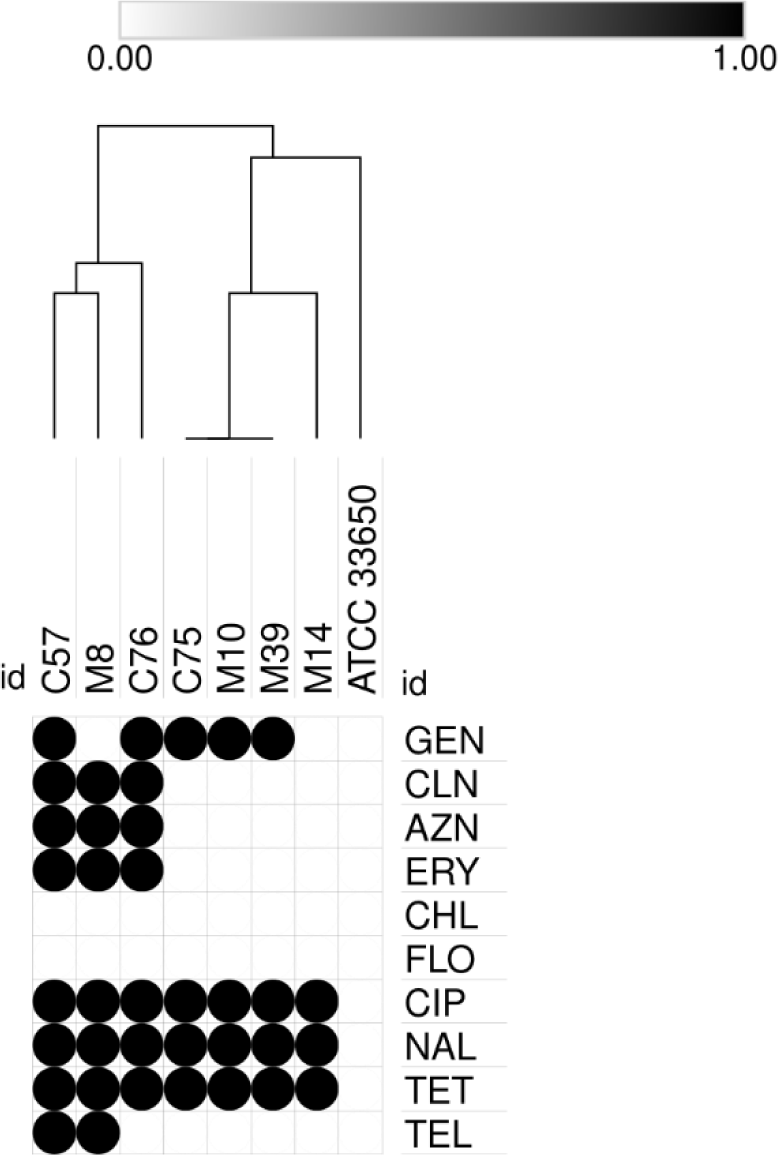
Antibiotic sensitivity testing for *C. coli*. *C. coli* from Vietnam, along with the control strain *C. jejuni* ATCC33650 were tested by microbroth dilution; MIC equal or greater than CLSI cut-off indicated by black circle. GEN (gentamicin), CLIN (clindamycin), AZN (azithromycin), ERY (erythromycin), CHL (chloramphenicol), FEN (florfenicol), CIP (ciprofloxacin), NAL (nalidixic acid), TET (tetracycline), TEL (telithromycin).

**Table 1.**
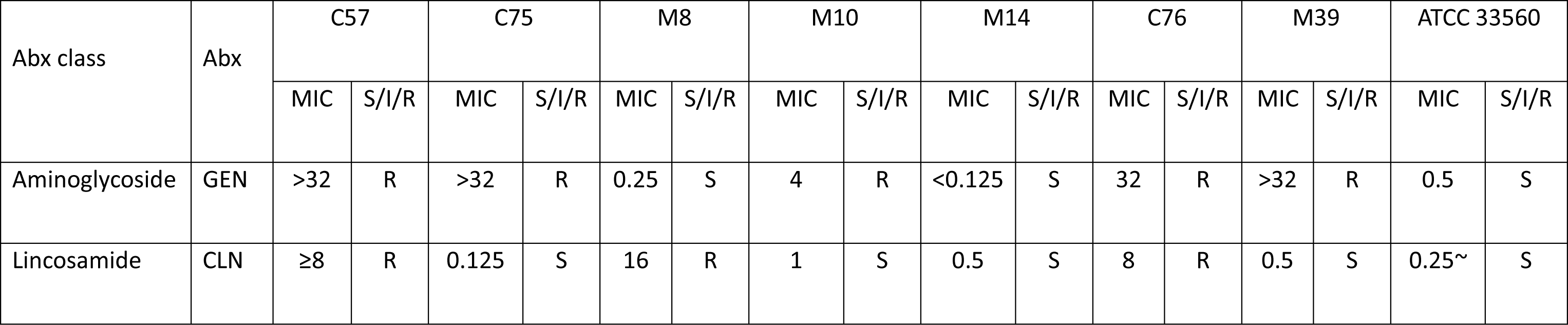
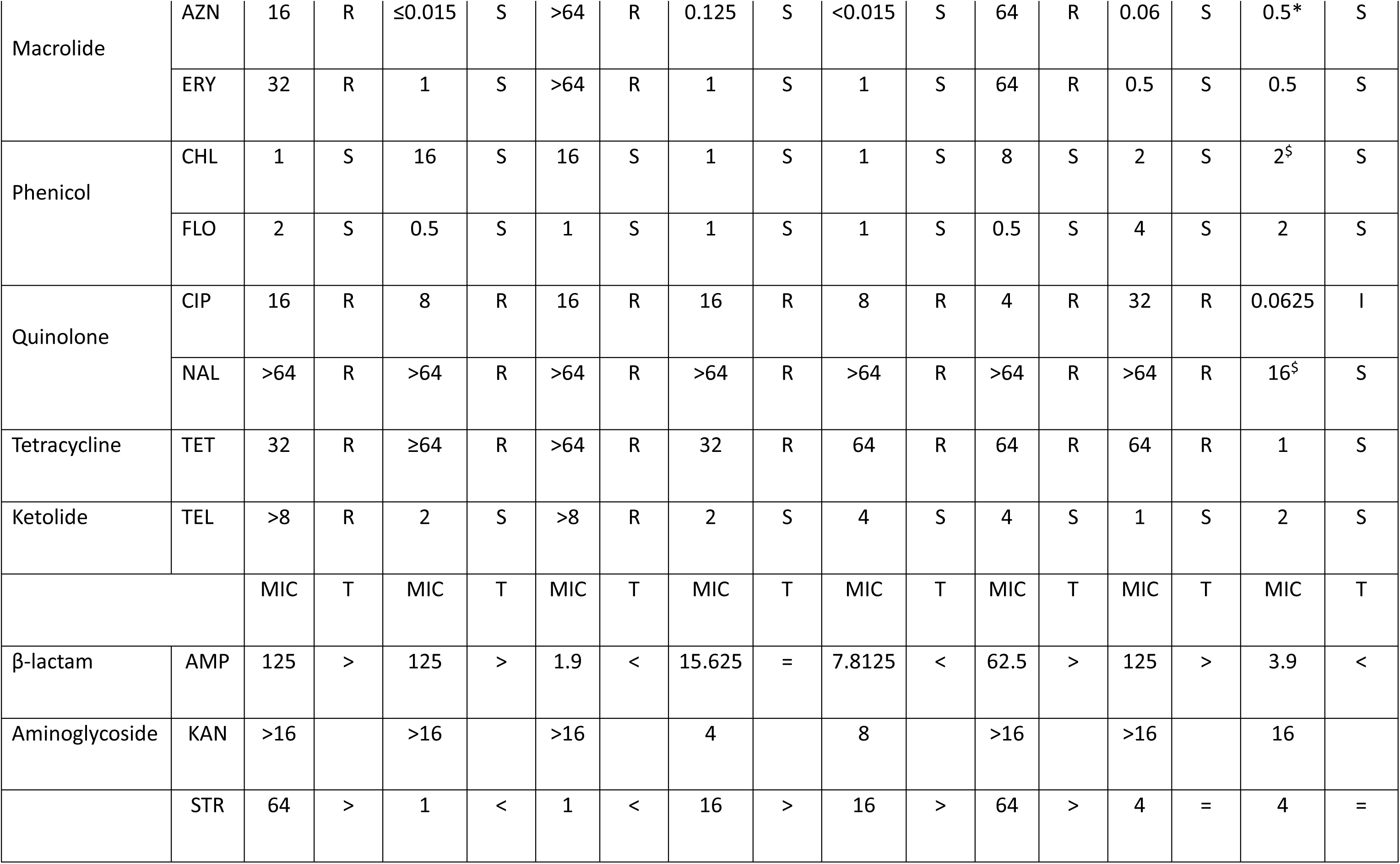
Antibiotic susceptibility of *C. coli* isolates. MIC values are in mg/L for *C. coli* strains C57, C75, M8, M10, M14, C76, M39 and *C. jejuni* ATCC 33560. S (susceptible), I (intermediate), R (resistant), Abx (antibiotics tested), GEN (gentamycin), CLN (clindamycin), AZN (azithromycin), ERY (erythromycin), CHL (chloramphenicol), FLO (florfenicol), CIP (ciprofloxacin), NAL (nalidixic acid), TET (tetracycline), TEL (telithromycin), AMP (ampicillin), KAN (kanamycin), STR (streptomycin). T (tentative epidemiological cut-off values [(T)ECOFF]); AMP [16 mg/L] and STR [4 mg/L], KAN has no EUCAST or CLSI breakpoint or (T)ECOFF value for *Campylobacter* strain*s*, > (greater than (T)ECOFF), = (equivalent to (T)ECOFF, < (less than (T)ECOFF). *C. jejuni* ATCC 33560 within expected MIC range unless above expected MIC (*), below expected MIC (∼) or no given MIC ($).

#### 3.1.4. Galleria mellonella pathogenicity

*Campylobacter* does not naturally infect *G. mellonella*, however, due to the lack of readily available models for testing *Campylobacter* virulence, it is an established surrogate infection model to assess virulence and it has been used for numerous bacterial infections, including *C. jejuni* (Kurstak and Vega 1968, Dudziak and Jozwik 1969, Champion et al. 2010, Insua et al. 2013, Viegas et al. 2013, Bojanic et al. 2020). Pathogenicity of *C. coli* isolates was tested using a *G. mellonella* wax moth infection model (Figure 3). The results showed that disease in *G. mellonella* infected with *C. coli* M10 and M14 are significantly attenuated (p=0.0438 and p=0.0290 respectively, Gehan-Breslow-Wilcoxon test) compared to the positive control *C. jejuni* 81-176, with many *G. mellonella* surviving beyond 9 days. Survival of *G. mellonella,* infected with *C. coli* C75, C76, M8 and M39, was not significantly different from *C. jejuni* 81-176; p= 0.593, 0.117, 0.692 and 0.164 respectively (Figure 3). More *C. coli* C57 infected *G. mellonella* died than *C. jejuni* 81-176 over the initial 4 days, but this was not significant over the course of 9 days, p=0.224 (Figure 3).

**Figure 3.**
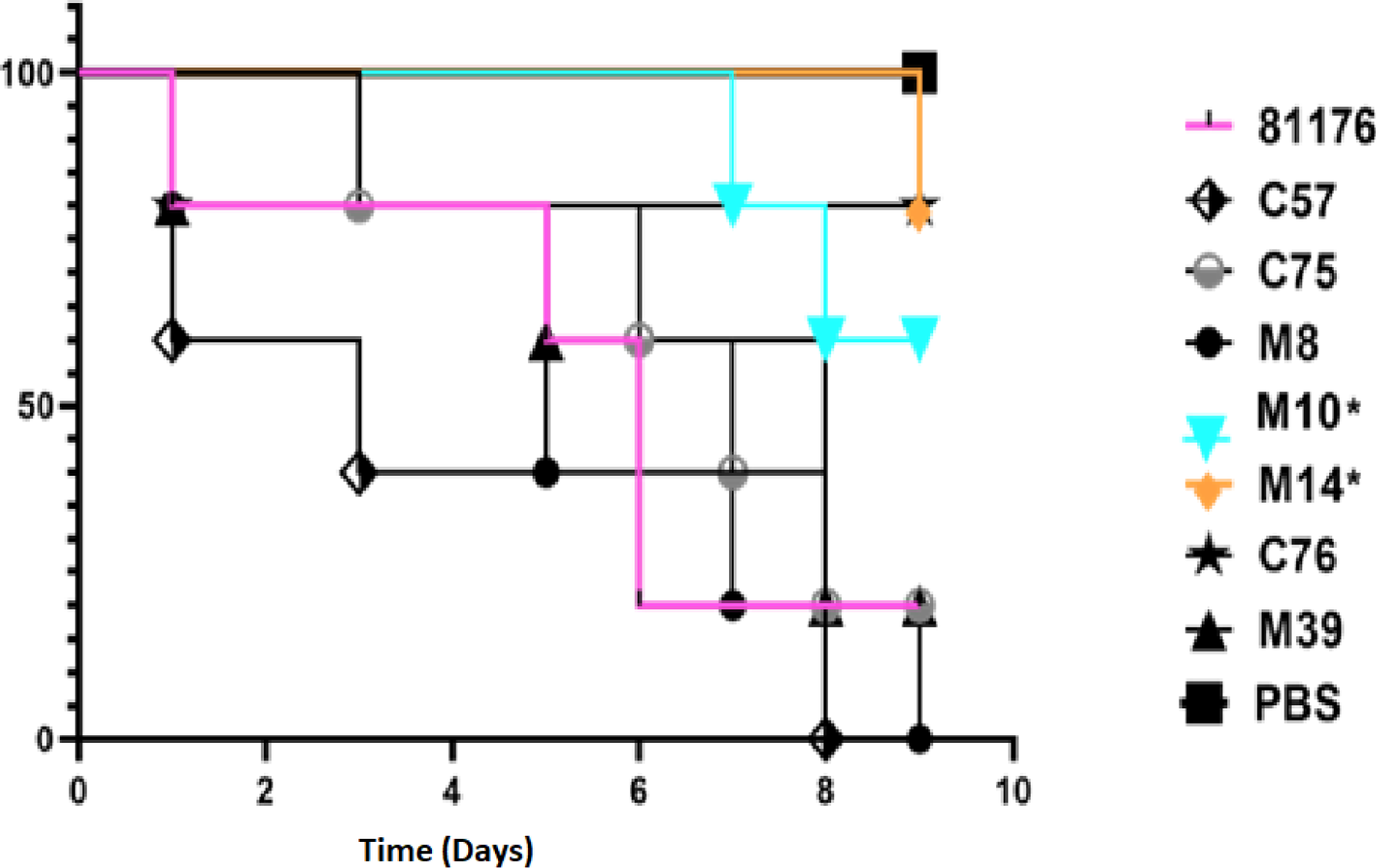
Virulence testing of Vietnamese *C. coli* isolates using *Galleria mellonella* model. Representative survival plot showing probability of survival of *G. mellonella* after infection with *C. coli* isolates (C57, C75, M8, M10, M14, C76, M39). *C. jejuni* 81-176 was used as a positive reference standard and PBS negative control. Data from an independent biological repeat, representative of the three independent experiments conducted is shown. *C. coli* isolates M10 and M14 were significantly attenuated (p=0.0438 and p=0.0290 respectively, Gehan-Breslow-Wilcoxon test) compared to the *C. jejuni* 81-176 reference control.

**Figure 4.**
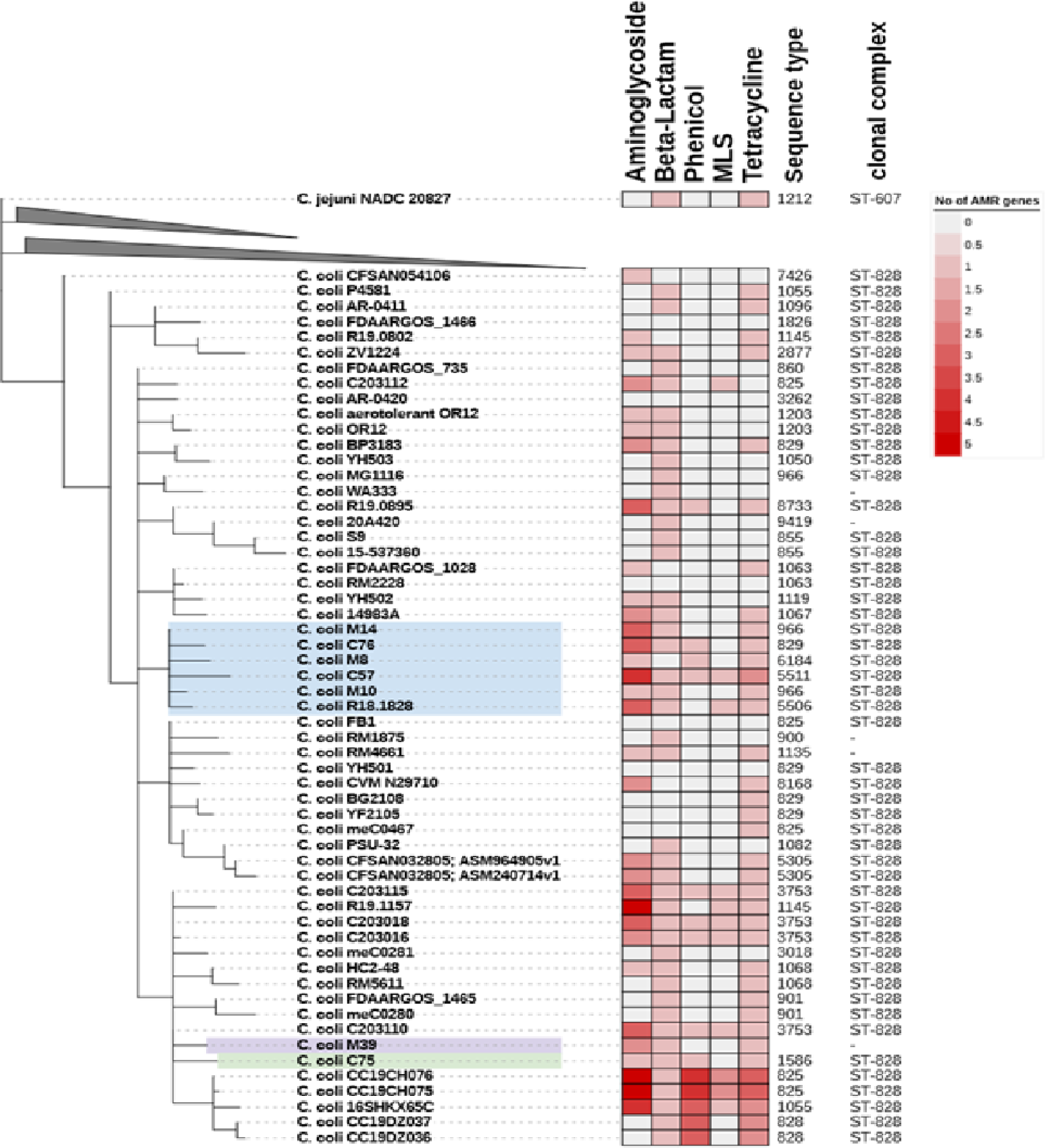
Phylogenetic analysis of clonal complex 828 (CC ST-828) isolates. Core-genome MLST phylogeny for seven Vietnamese *C. coli* isolates against RefSeq complete *C. jejuni* and *C. coli* isolates. Only *C. coli* isolates are shown, all of which have CC ST-828, as determined by PubMLST database. chewBACCA with schema having 2794 loci and 299214 alleles was used to determine cgMLST. cgMLST output was used to build phylogenetic tree on interactive tree of life (iTOL). *C. jejuni* NADC 20827 (ST1212, CC ST-607) was used as outlier to root phylogeny. Genome analysis using Abricate and ResFinder database identified between 0 and 5 resistance genes linked to five antibiotic classes (heatmap). MLS (Macrolides, Lincosamides, Streptogramins).

### 3.2 Genotypic analysis

#### 3.2.1. *C. coli* genomes

All isolates were whole genome sequenced using Illumina short read sequencing technology with *C. coli* isolates C57, C76, M8 and M39 further sequenced using a Minion sequencer (Supplementary Table A2). Draft genomes ranged from 1.64 – 1.77 Mbp, which is comparable to *C. coli* 76339 accession HG326877 (1.58 Mbp) and *C. coli* RM4661 accession CP007181 (1.82 Mbp). Assembly contig numbers ranged from 1 (M8) to 21 (C75) contigs of length ≥ 1kb. The mean number of predicted coding sequences (CDSs) for sequenced *C. coli* strains was 1773 (SD ±50.2) (Supplementary Table A2).

#### 3.2.2. Identification of a novel *C. coli* plasmid

A circularised plasmid of length 27,557 bp was identified for isolate *C. coli* M39, named pCCM39-Huong. Annotation of pCCM39-Huong identified 37 coding sequences; 22 had predicted function, but 15 (40%) were hypothetical of unknown function (Supplementary Table A3). Annotated genes included *ssb*2 (single stranded DNA binding protein) (Jeon and Zhang 2007), potentially involved in DNA processing during conjugation. Genes *traG* and *traL* both involved in conjugal transfer of genetic material, a toxin component (*vbhT*) (Sprenger et al. 2017), *virB1*, *virB4*, *virC, trbB*, *traM*, *trbl*, *trbF*, *trbF*, *trbM* and *trbD* all related to type IV secretion system were also found (Zatyka and Thomas 1998, Mace et al. 2022) (Supplementary Table A3).

pCCM39-Huong had sequence similarity to two unnamed plasmids found in *C. coli* isolates 20A420 (accession CP058341) and *C. coli* meC0281 (CP027635) (Figure 5). *C. coli* 20A420 plasmid was of length 26,486 bp, with 97.36% identity and 90 % coverage compared to pCCM39-Huong. Plasmid pCCM39-Huong contained an additional three non-contiguous hypothetical proteins compared to p20A420 (Figure 5). Plasmid meC0281 was of length 30,719 bp and had 96.33% identity and 96% coverage when compared to pCCM39-Huong (Figure 5). Plasmid pCCM39-Huong contained an additional two non-contiguous hypothetical proteins compared to pmeC0281 (Figure 5). *C. jejuni* 81-176 contains two plasmids (pTet [AF226280] and pVir [AY394561]) both of approximately 35kb. pCCM39-Huong reads did not map to any of the pVir gene sequence, demonstrating that these plasmids are highly different to plasmid pCCM39-Huong. Comparison between pCCM39-Huong assembly and pTet only identified a single shared complete gene (a putative DNA primase) and partial matches to *cpp32* (phage/Rha family transcriptional regulator) and *cmgB5* (putative type IV secretion system component). Mapping of the M39 genome fastq data showed that pTet *tetO* (tetracycline resistance) was highly similar to *tetO* present within the M39 genome.

**Figure 5.**
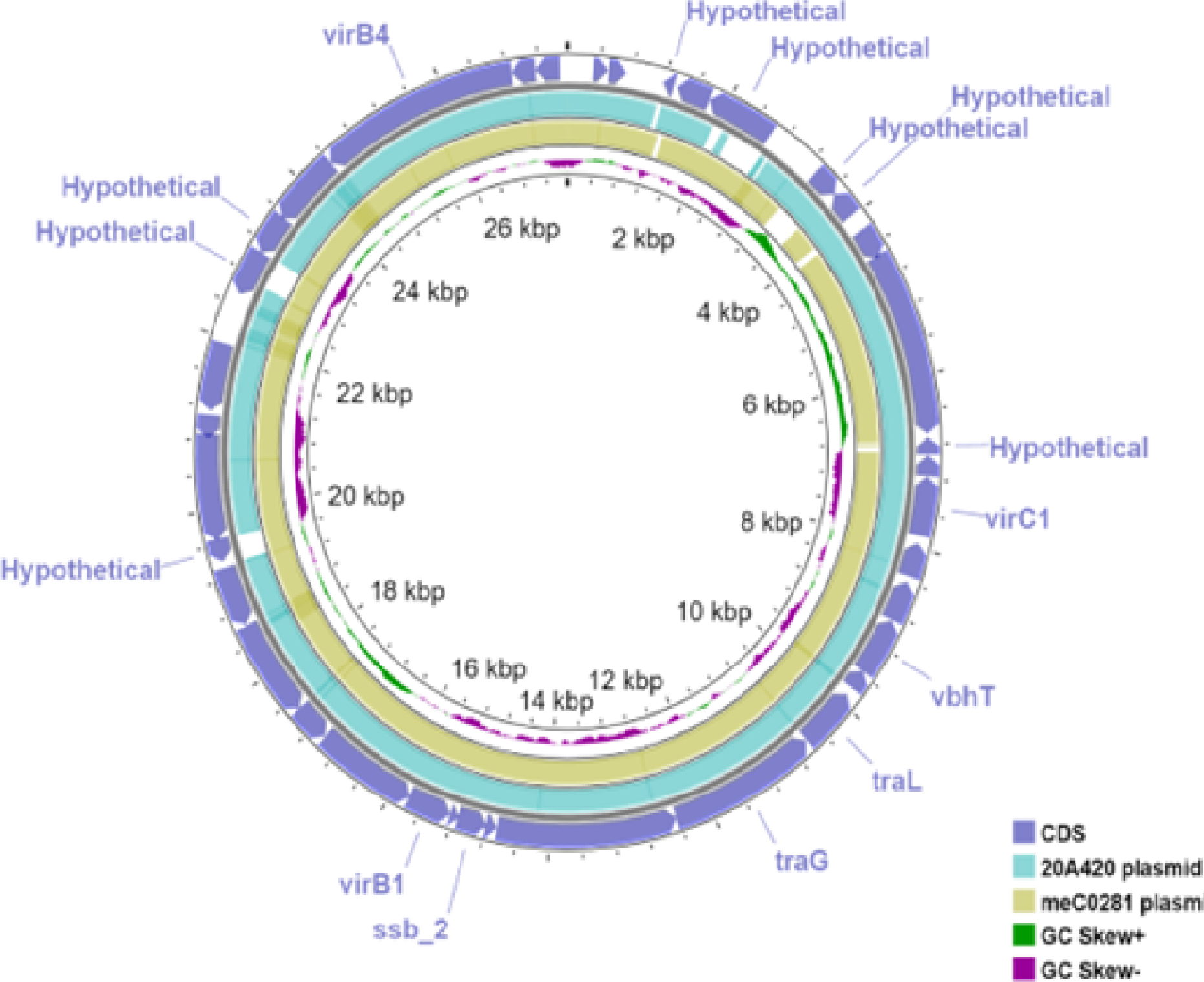
*C. coli* plasmid pCCM39-Huong. A circular image representation of pCCM39-Huong generated using CG view, showing GC skew, blast alignment to two closest plasmid sequences plasmid 20A420 and meC0281. CDSs are left annotated where genes were not hypothetical or where there are sequence differences between pCCM39-Huong and plasmid 20A420 and meC0281.

#### 3.2.3 *C. coli* epidemiological information

Analysis of the sequenced *C. coli* genomes identified five different sequence types (STs), with M39 having a novel ST, due to a novel *glnA* allele that was one single-nucleotide polymorphism (T198C) different to *glnA*(291) (Table 2). All STs belonged to the ST-828 clonal complex (Table 2). In the PubMLST database (accessed 22-6-2022) of the 16,083 ST-828 clonal complex isolates, 15,974 (99.3 %) were identified as *C. coli*, 97 *C. jejuni* and 12 as *Campylobacter* spp. cgMLST further confirmed that the isolates were *C. coli*, as these were grouped away from *C. jejuni* (Figure 4) and clustered with reference ST-828 clonal complex isolates (Figure 4). The sequenced isolates were not clustered based on whether they were isolated from farm or supermarket/retail market (Figure 4). *C. coli* M39 and C75 belonged to the same subclade (Figure 4) and shared 5 MLST alleles (Table 2). M10 and M14 (from supermarket-bought chicken meat) were both ST-966, however cgMLST data indicated that these were very similar, but not identical isolates. *C. coli* M14, C76, M8, C57, M10 belonged to the same subclade and were related to *C. coli* R18.1828, isolated from Taiwan (Figure 4).

**Table 2.** Multi-locus sequence type (MLST) profile for *C. coli* isolates. MLST data was extracted from draft genome sequences using the PubMLST *Campylobacter jejuni/coli* database. ST (sequence type), CC (clonal complex). * M39 had a novel *glnA* allele that was one SNP different to the *glnA* (291) allele.

| Isolate | Source | ST<br>(CC) | Allele |  |  |  |  |  |  |
| --- | --- | --- | --- | --- | --- | --- | --- | --- | --- |
|  |  |  | *aspA* | *glnA* | *gltA* | *glyA* | *pgm* | *tkt* | *uncA* |
| C57 | Chicken faeces<br>(Farm) | ST5511<br>(ST-828) | 33 | 39 | 30 | 82 | 113 | 47 | 139 |
| C75 | Chicken faeces<br>(Farm) | ST1586<br>(ST-828) | 33 | 176 | 30 | 82 | 113 | 43 | 17 |
| M8 | Chicken meat<br>(Retail Market) | ST6184<br>(ST-828) | 32 | 39 | 30 | 82 | 113 | 43 | 17 |
| M10 | Chicken meat<br>(Supermarket) | ST966<br>(ST-828) | 33 | 39 | 30 | 79 | 112 | 47 | 17 |
| M14 | Chicken meat<br>(Supermarket) | ST966<br>(ST-828) | 33 | 39 | 30 | 79 | 112 | 47 | 17 |
| C76 | Chicken faeces | ST829 | 33 | 39 | 30 | 82 | 113 | 43 | 17 |
|  | (Farm) | (ST-828) |  |  |  |  |  |  |  |
| M39 | Chicken meat<br>(Retail market) | -<br>(ST-828) | 33 | 291* | 30 | 82 | 113 | 47 | 17 |

ST-5511 (*C. coli* C57) had previously been described in three *C. coli* isolates from China, with two identified from chicken. ST-1586 (*C. coli* C75) has been identified in 45 isolates including six from North America, five from the USA, and ten from the UK, all of which were identified from human faeces, and some documented as a sporadic case of gastroenteritis. ST-1586 (*C. coli* C75) has been reported from Vietnamese clinical samples between 2014-2019 and also from China, USA, Canada, UK, Spain, Germany, and other European countries. *C. coli* ST-6184 (*C. coli* M8) has been identified previously in poultry in the USA and China, with a single case from China isolated from human stool, linked to an outbreak. ST-966 (M14 and M10) has mostly been documented from the USA, except a single UK clinical isolate. ST-829 (*C. coli* C76) has been submitted over 1200 times to the MLST database, mostly identified in high-income countries (HICs), mainly from the USA, with 1021 isolates representing 83% of the documented cases. Forty-four cases of ST-829 was identified in England and 33 from Scotland, all of which were isolated from human faeces, relating to gastroenteritis.

#### 3.2.4. AMR genotypes

Analysis of the draft genomes identified potential AMR genes to all classes of antibiotics tested. The β-lactamase gene *bla*_OXA-193_ was present in all sequenced genomes except M8 and 142 out of 284 (50%) reference genomes (Supplementary Table A4). Four aminoglycoside genotypes were identified with isolates containing between one (M8, M10, C75) and three (C57, C76, M14) CDSs (Supplementary Table A4). All isolates had a copy of *tet* (O) linked to tetracycline resistance, 4 out of 7 sequenced isolates had *cat*2 phenicol resistance gene, but only C57 had the *erm*(B) MLS (macrolide, lincosamide and streptogramin) resistance gene (Figure 4, Supplementary data S4).

Sequence analysis showed that only *C. coli* C57 was predicted to have nine resistance genes, covering aminoglycosides, β-lactam, phenicol, MLS and tetracycline resistance (Figure 4, Supplementary Table A4). Analysing the *C. coli* M8 genome identified aminoglycosides, β-lactamase and phenicol AMR genes (Figure 4, Supplementary Table A4). SNP analysis identified ciprofloxacin resistance due to a gyrase subunit A (*gyrA)* (T86I) mutation in all isolates. The ResFinder database did not identify any resistance genes for the control sensitive strain *C. jejuni* ATCC 33560.

### 3.3 Correlation between AMR phenotype and genotypes

Prediction of phenotypic resistance to antibiotics from the presence of genes linked to resistance has proven complex (Doyle et al. 2020). All isolates, except M8 and M14 were resistant to gentamycin. Four isolates had an MIC above streptomycin (T)ECOFF and five had an MIC of >16 mg/L for kanamycin, however all isolates contained at least one aminoglycoside resistance gene. Only the isolates (*C. coli* C57, M10, M14 and C76) with *ant*(6)-Ia were resistant to streptomycin. *C. coli* M8 possessed only *aph*(3’)-III and had a kanamycin MIC > 16, similar to other three other *aph*(3’)-III positive isolate (*C. coli* C75, C76 and M39). The fifth *aph*(3’)-III positive isolates, *C. coli* M14, had a kanamycin MIC of 8. While C57 did not possess an *aph*(3’)-III and had a kanamycin MIC >16 this isolate did have four alternative aminoglycoside genes. All isolates, except *C. coli* M8 possessed *bla*_OXA-193_ as the only β-lactamase yet MIC values ranged from 1.9 (*C. coli* M8) to >125 (*C. coli* C57, C75 & M39). *C. coli* C57, M8 and C76 were resistant to erythromycin, however the *erm*(B) inducible erythromycin resistance was only present in *C. coli* C57.

#### 3.4 3 Correlation between virulence related genes and genotypes

Genotypic data for each isolate showed that motility genes (*flaA*, *flaB*, *flaC*, *flaG*, *flgB*, *flgC*, *flgD*, *flgE2*, *flgE*, *flgG2*, *flgG*, *flgH*, *flgI*, *flgK*, *flgL*, *flgR*, *flhA*, *flhB*, *flhF*, *flhG*, *fliA*, *fliD*, *fliE*, *fliF*, *fliG*, *fliH*, *fliI*, *fliL*, *fliM*, *fliN*, *fliP*, *fliQ*, *fliR*, *fliS*, *fliY*, *motA*, *motB*, *pflA*) were present in all *C. coli* isolates. All *C. coli* isolates also had the adhesion (*cadF*, *jlpA*, MOMP, PEB1) (Krause-Gruszczynska et al. 2007), cytolethal toxin *(cdtABC)* and *Campylobacter* invasion related genes (*ciaB*) (Konkel et al. 1999). The well-characterised *O-* and *N*-linked general glycosylation systems were also present. Surprisingly, *C. coli* C57, C75, M10 and M14 also possessed *cysC1* commonly found in a plant pathogen *Pseudomonas syringae,* encoding for a phytotoxin phaseolotoxin (Arrebola et al. 2011).

*C. coli* M10 (and M14) demonstrated significantly lower virulence in the *G. mellonella* infection model and was the least motile (along with M8), whereas *C. coli* C57 was the most virulent isolate. To identify any genetic cause for the virulence of *C. coli* C57, we isolated C57 fastq reads that did not match the M10 and M14 assembled draft genomes. These unmapped reads were assembled to represent CDSs in *C. coli* C57 that were absent or highly divergent from *C. coli* M10 and M14. The unmapped reads assembled in to 39 genome fragments containing 158 predicted CDSs. To remove any CDSs that were present in M10 and M14, but were not currently in these draft genomes, the raw reads from M10 and M14 were mapped against this assembly, which identified six contigs containing 16 CDSs that were present in M14 but not M10. 106 of 142 CDS were annotated as hypothetical proteins, two were IS1595 family transposases, 28 were annotated as enzymes, with many linked to sugar metabolism (e.g. dTDP-glucose 4,6-dehydratase), modification methylase (*dpnIIA* and *dpnIIB*), ribosomal silencing factor (RsfS) and sugar efflux transporter. Additionally, both flagellin FlaA (highly like the residues 102-572 of 11168 FlaA) and FlaB (67 amino acid fragment, identical to last 67 residues of 11168 FlaB) were present, demonstrating significant variation between *C. coli* C57 and M10/M14 flagellin genes.

## 4. Discussion

Infectious disease profiles can change over time, particularly in low-income regions, where evolving economic conditions may prompt changes in human behaviour and in veterinary practices. These changes highlight the need for continuous monitoring of foodborne pathogens to build comprehensive virulence profiles. *C. coli,* despite being less studied than *C. jejuni,* has emerged as a significant public health concern due to its increasingly alarming profile of antimicrobial resistance, often leading to multi-drug resistance(van Vliet et al. 2022). Additionally, there is evidence of genetic convergence between *C. coli* and *C. jejuni*, due to human activity, such as intensive farming of poultry, where both species can inhabit, allowing increased horizontal transmission of DNA between the species (Sheppard et al. 2008). Seven *C. coli* isolates from the Vietnamese food chain were isolated and fully characterised. The MLST profiles of *C. coli* C75 (ST-1586), M8 (ST-6184), M10 (ST-966) and M14 (ST-966) have previously been found in human stool samples, many of which were linked to sporadic cases of gastroenteritis, demonstrating that the isolates could potentially be of clinical importance. All the MLST profiles, apart from M39, were identified in HICs, with all 32 ST-966 isolates in the MLST database (25^th^ Jan 2022) being from HICs, along with 1021 (83% of isolates) isolates for ST-829 (*C. coli* C76) from the USA, many of which from carriers such as poultry.

The antibiotic of choice for *Campylobacter* associated gastroenteritis are macrolides, although fluoroquinolones are also commonly used (Wieczorek and Osek 2013), however three out of seven isolates were found to be resistant to both macrolides in this study. The resistance profile of *Campylobacter* to macrolide tends to be continent specific, however macrolide resistance is found to be more common in *C. coli* isolates than *C. jejuni* (Luangtongkum et al. 2009). Macrolides resistance in North America is around 10% for both *C. jejuni* and *C. coli* isolates isolated from humans/broilers. Yet in Europe, a far larger number of *C. coli* isolates are macrolide resistant, when compared to *C. jejuni* (Eurosurveillance editorial 2014). While in Asia, less than 5% of *C. jejuni* were macrolide resistant, as opposed to 14 – 62% of *C. coli* (Luangtongkum et al. 2009). Fluoroquinolones resistance (ciprofloxacin) present in all isolates was through a *gyrA* mutation, that has been shown to be linked with ciprofloxacin resistance (Ohishi et al. 2017, Devi et al. 2019). This was unsurprising, as previous studies have indicated *gyrA* related *Campylobacter* spp. resistance to fluoroquinolones being common in Vietnam, with *Campylobacter* isolates rapidly acquiring resistance (Luangtongkum et al. 2009). Tetracycline resistance was also observed for all isolates due to *tet(O)* (Pratt and Korolik 2005), although a duplicate copy of *tet(O)* in *C. coli* C57 was not linked with higher MIC. Tetracycline is seldom used for the treatment of acute gastroenteritis (Garcia-Fernandez et al. 2018), but is the most frequent antibiotic in the treatment and control of various poultry diseases, leading to high resistance rates in a number of zoonotic pathogens. All *C. coli* isolates, apart from M8 possessed a *bla*_OXA-193_, which was also present also in 142 reference genomes, and most of these isolates had an ampicillin MIC equal to or far higher than the (T)ECOFF. The sensitivity to ampicillin in M8 was unexpected with the presence of *bla*_OXA-193_ but the reason for the lack of expression was not clear from the draft genome.

The (T)ECOFF was determined by analysis of 419 *C. coli* identified between January 2014 and September 2015 in France, with only 7 out of 114 (6.1%) having an MIC of 128 mg/L or higher however 3 out of 7 (43%) of Vietnamese isolates had an MIC of 125 mg/L (Sifre et al. 2015). The presence of multiple AMR phenotypes meant that all, but *C. coli* M14 were resistant to at least three classes of antibiotics, however no isolates demonstrated resistance to the phenicols tested. Florfenicol has the same mechanism of action as chloramphenicol and is used for agricultural purposes. Florfenicol is also thought to be more active than chloramphenicol (Sutili and Gressler 2021). Yet, the borderline resistance for chloramphenicol in two of the *C. coli* strains C75 and M8 should be concerning, as resistance to this group of antibiotics may arise without judicious use.

In this study we characterised the phenotypic and genotypic profile of *C. coli* isolates and investigated their virulence potential, compared to the more commonly studied *C. jejuni* species. Genome sequence analysis identified common *Campylobacter* sp. virulence related genes with most isolates equally motile as the reference hyper-motile *C. jejuni* 11168H strain and equally lethal as the reference strain *C. jejuni* 81-176 in the *G. mellonella* model of infection. Genome sequencing identified a potential virulence related plasmid, carrying genes related to the type IV secretion system. The type IV secretion system is involved in DNA conjugation, helping bacteria in adapting to environmental stress and is a potential mechanism of spreading AMR, along with transformation, allowing the uptake of DNA from the environment (Wallden et al. 2010). The type IV secretion system can also play a role in increased host colonisation, biofilm formation and transfer of toxins to host cells and thus can have a direct involvement in virulence (Kienesberger et al. 2011).

Although *C. jejuni* is the most common *Campylobacter* species to infect humans, other *Campylobacter* species, such as *C. coli* need to be studied for several reasons, including interspecies horizontal gene transfer (Mourkas et al. 2022), which may contribute the emergence of multi-drug resistant pathogens. This, together with the expanding global poultry production networks and the intensification of production, may pose a global public health risk. This highlights the urgent need for a coordinated global effort to combat AMR.

## 5. Authors and contributors

BL, LQH, RAS Conceptualization. BL, GN, AC, FN, SW data curation and investigation. BL, GN, SW, RAS Formal analysis and methodology. RAS Funding acquisition and project administration. BL, GN, BWW, RAS Writing – original draft. All authors Writing – review & editing.

## 7. Declaration of generative AI in scientific writing

During the preparation of this work the authors did not use any generative AI and AI-assisted technologies in the writing process.

## 9. Ethical approval

Not required

## 10. Consent for publication

Not required

## 11. Acknowledgements

We would like to thank Abdi Elmi and Sherif Abouelhadid for their technical assistance.

## 14. Supplementary files

**Table A1. Identification of Vietnamese *Campylobacter* isolates from Hanoi, Vietnam.** Phenotypic tests used to identify *Campylobacter* genus and species. Results are consistent with *C. coli*. – (negative), + (positive).

**Table A2. Genome assembly summary.** All *C. coli* isolates were sequenced with Illumina paired-end short read technology. *C. coli* C57, C76, M8 and M39 were additionally sequenced with MinION long-read technology. \**C. coli* M39 assembly statistics includes pCCM39-Huong plasmid of length 27,557 bp. CDS (Coding sequences).

**Table A3. *C. coli* M39 plasmid predicted proteins.** Automatic annotation indicates Prokka automated annotation. BLASTP indicates additional annotation obtained using protein-protein BLAST against the NCBI non-redundant database. Percent similarity indicates amino acid match to the given db_xref protein.

**Table A4. Analysis of the draft genomes identified potential AMR genes**. Abricate with the ResFinder database (update date 10-03-2022) was used to identify AMR genes.

