## Supplementary information for "Understanding *Campylobacter coli* isolates from the Vietnamese meat production network; a pilot study"

**Supplementary data**

**Table S1**. **Identification of Vietnamese *Campylobacter* isolates from Hanoi, Vietnam.** Phenotypic tests used to identify campylobacter genus and species. Results are consistent with *C. coli*. – (negative), + (positive).

| Isolate | Source | Location | Growth at 25 °C | Morphology | Catalase | Oxidase | Indole | Hippurate |
|---|---|---|---|---|---|---|---|---|
| C57 | Chicken faeces | Farm | - | Campy-typical, spiral | + | + | - | - |
| C75 | Chicken faeces | Farm | - | Campy-typical, spiral | + | + | - | - |
| C76 | Chicken faeces | Farm | - | Campy-typical, spiral | + | + | - | - |
| M8 | Chicken meat | Retail market | - | Campy-typical, spiral | + | + | - | - |
| M10 | Chicken meat | Supermarket | - | Campy-typical, spiral | + | + | - | - |
| M14 | Chicken meat | Supermarket | - | Campy-typical, spiral | + | + | - | - |
| M39 | Chicken meat | Retail market | - | Campy-typical, spiral | + | + | - | - |

**Table S2. Genome assembly summary**. All *C. coli* isolates were sequenced with Illumina paired-end short read technology and with *C. coli* C57, C76, M8 and M39 additionally sequenced with MinION long-read technology. **C. coli* M39 assembly statistics includes pCCM39-Huong plasmid of length 27,557 bp. CDS (Coding sequences).

| **Isolate** | **C57** | **C75** | **M8** | **M10** | **M14** | **C76** | **M39*** | **Mean** | **SD** |
|---|---|---|---|---|---|---|---|---|---|
| Total Length (bp) | 1,710,747 | 1,771,751 | 1,769,631 | 1,725,657 | 1,732,229 | 1,640,312 | 1,724,504 | 1,724,976 | 43,981 |
| Largest Contig (bp) | 1,655,035 | 383,932 | 1,769,631 | 783,471 | 925,559 | 1,220,521 | 672,432 | 1,058,654 | 514,007 |
| N50 (bp) | 1,655,035 | 264,610 | 1,769,631 | 462,807 | 925,559 | 1,220,521 | 531,949 | 975,730 | 594,404 |
| Contigs | 9 | 21 | 1 | 19 | 16 | 5 | 11 | 12 | 7.36 |
| GC | 31.5 % | 31.4 % | 31.4 % | 31.4 % | 31.4 % | 31.5 % | 31.4 % | 31.4 % | 0.0005 |
| Total genes | 1,780 | 1894 | 1861 | 1823 | 1838 | 1754 | 1839 | 1,827 | 47.4 |
| CDS | 1,723 | 1848 | 1805 | 1767 | 1782 | 1698 | 1788 | 1,773 | 50.2 |
| rRNA | 10 | 3 | 9 | 9 | 9 | 9 | 6 | 8 | 2.5 |
| tRNA | 44 | 40 | 44 | 44 | 44 | 44 | 42 | 43 | 1.6 |

**Table S3. *C. coli* M39 plasmid predicted proteins.** Automatic annotation indicates Prokka automated annotation. BLASTP indicates additional annotation obtained using protein-protein BLAST against the NCBI non-redundant database. Percent similarity indicates amino acid match to the given db_xref protein.

| Systematic ID | Automatic annotation | Length | BLASTP matches | Similarity | db_xref |
|---|---|---|---|---|---|
| pCCM39-001 | hypothetical | 174 | P-type conjugative transfer ATPase TrbB | 100% | GI:1098456436 |
| pCCM39-002 | hypothetical | 204 | TrbB | 97.24 | GI:2193642593 |
| pCCM39-003 | hypothetical | 135 | DNA-binding protein | 98.64% | GI:1444891879 |
| pCCM39-004 | hypothetical | 420 | | | |
| pCCM39-005 | hypothetical | 843 | | | |
| pCCM39-006 | hypothetical | 336 | | | |
| pCCM39-007 | hypothetical | 324 | | | |
| pCCM39-008 | hypothetical | 384 | DNA transfer protein | 100% | GI:1098456441 |
| pCCM39-009 | hypothetical | 2,157 | Relaxase/mobilization nuclease domain-containing protein | 98.86% | GI:2093938767 |
| pCCM39-010 | hypothetical | 201 | | | |
| pCCM39-011 | hypothetical | 240 | | | |
| pCCM39-012 | virC1 | 687 | AAA family ATPase CDS | 100% | GI:2093938769 |
| pCCM39-013 | hypothetical | 396 | Conjugal transfer protein TraM | 100% | GI:2093938770 |
| pCCM39-014 | hypothetical | 468 | LepB | 100% | GI:2093938771 |
| pCCM39-015 | vbhT | 684 | | | |
| pCCM39-016 | hypothetical | 243 | | | |
| pCCM39-017 | traL | 720 | Conjugal transfer protein TraL CDS | 100% | GI:2093938774 |
| pCCM39-018 | traG | 1,824 | | | |
| pCCM39-019 | hypothetical | 2,214 | ssDNA-binding domain-containing protein | 98.50% | GI:2093938776 |
| pCCM39-020 | hypothetical | 141 | | | |
| pCCM39-021 | ssb 2 | 363 | | | |
| pCCM39-022 | hypothetical | 102 | Ssb | 100% | GI:2076397817 |
| pCCM39-023 | virB1 | 546 | | | |
| pCCM39-024 | hypothetical | 1,242 | Conjugal transfer protein TrbI | 100% | GI:1780455738 |
| pCCM39-025 | hypothetical | 444 | | | |
| pCCM39-026 | hypothetical | 1,155 | TrbG | 100% | GI:2093938753 |
| pCCM39-027 | hypothetical | 696 | Conjugal transfer protein TrbF | 99.84% | GI:2093938754 |
| pCCM39-028 | hypothetical | 270 | | | |

| | | | | | |
|---|---|---|---|---|---|
| pCCM39-029 | hypothetical | 1,224 | | | |
| pCCM39-030 | hypothetical | 270 | | | |
| pCCM39-031 | hypothetical | 795 | | | |
| pCCM39-032 | hypothetical | 555 | | | |
| pCCM39-033 | hypothetical | 483 | | | |
| pCCM39-034 | hypothetical | 867 | TrbM | 97.44% | GI:769549900 |
| pCCM39-035 | virB4 | 2,466 | | | |
| pCCM39-036 | hypothetical | 288 | Conjugal transfer protein TrbD | 100% | GI:1774382551 |
| pCCM39-037 | hypothetical | 297 | TrbC/VirB2 family protein | 100% | GI:2093938761 |

**s**

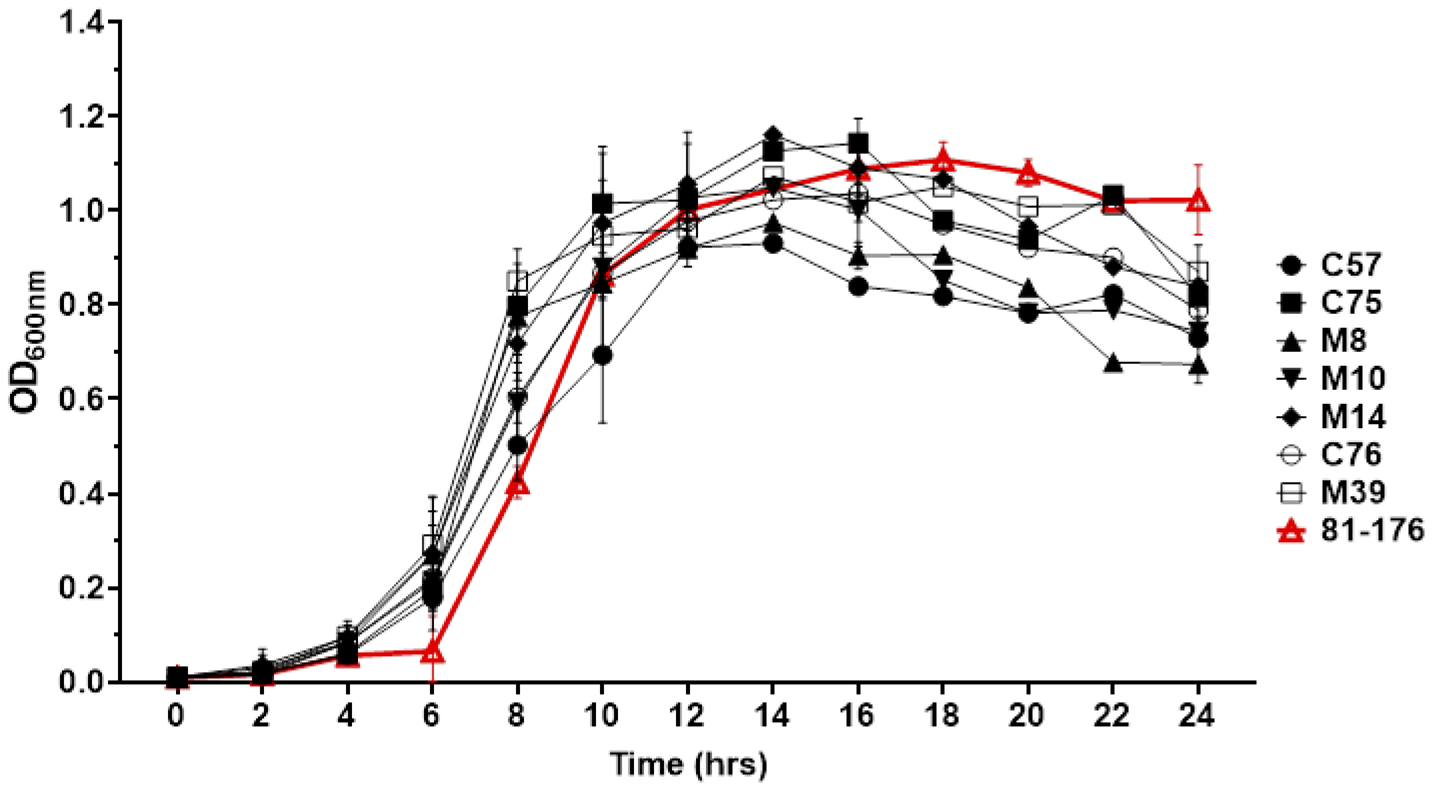


**Figure S1: Growth curve of *C. coli* isolates.** The isolates are represented as; *C. coli* C57 (circle), C75 (square), M8 (triangle), M10 (Nabla), M14 (Diamond), C76 (open circle), M39 (open square) and 81-176 (Red triangle) each grown in brucella broth**.** Bacteria were initially grown for 24 hrs on CBA and were suspended in 1 ml brucella broth. $OD_{600nm}$ was adjusted to 0.1 (~$10^8$ cfu) in 10 ml brucella broth. Growth was monitored by $OD_{600nm}$ measurements at 2 hrs intervals for 24 hrs at 37°C in a microaerobic chamber (Don Whitley Scientific, UK) containing 85% N2, 10% CO2, and 5% O2. *C. jejuni* strain 81-176 was used as control.
